# Single cell RNA-seq analysis reveals compartment-specific heterogeneity and plasticity of microglia

**DOI:** 10.1101/2020.03.10.985895

**Authors:** Junying Zheng, Wenjuan Ru, Jay R Adolacion, Michael S Spurgat, Xin Liu, Subo Yuan, Rommel X Liang, Jianli Dong, Andrew S Potter, S Steven Potter, Navin Varadarajan, Shao-Jun Tang

**Author notes:** These authors contributed equally to this work (Junying Zheng, Wenjuan Ru, Jay R Adolacion). Corresponding author: Shao-Jun Tang, Department of Neuroscience, Cell Biology, & Anatomy, The University of Texas Medical Branch, 301 University Blvd, Galveston, TX.

## Abstract

Microglia are heterogeneous and ubiquitous CNS-resident macrophages that maintain homeostasis of neural tissues and protect them from pathogen attacks. Yet, their differentiation in different compartments remains elusive. We performed single cell RNA-seq (scRNA-seq) analysis to compare the transcriptomes of microglia in adult mouse (C57/Bl) brains and spinal cords to identify microglial subtypes in these CNS compartments. Cortical microglia from 2-month mice consisted of a predominant population of the homeostatic subtype (HOM-M) and a small population (4%) of the inflammatory subtype (IFLAM-M), while spinal microglia consisted of 55% HOM-M and 45% IFLAM-M subtype. Comparison of cortical and spinal microglia at 2, 4 and 8 months revealed consistently a higher composition of the IFLAM-M subtype in the spinal cord. At 8-month, cortical microglia differentiated a small new subtype with interferon response phenotypes (INF-M), while spinal microglia polarized toward a proinflammatory phenotype, as indicated by the increase of microglia expressing IL-1β. To further characterize the differential plasticity of cortical and spinal microglial heterogeneity, we determined the microglial transcriptomes from HIV-1 gp 120 transgenic (Tg) mice, a model of HIV-associated neurological disorders. Compared with wild-type (Wt) cortical microglia, the gp120Tg cortical microglia had three new subtypes, with signatures of interferon I response (INF-M), cell proliferation (PLF-M), and myelination or demyelination (MYE-M) respectively; while INF-M and PLF-M subtypes presented at all ages, the MYE-M only at 4-month. In contrast, only the INF-M subtype was observed in the spinal microglia from 2- and 4-month gp120tg mice. Bioinformatic analysis of regulated molecular pathways of individual microglial subtypes indicated that gp120 more severely impaired the biological function of microglia in cortices than in the spinal cord. The results collectively reveal differential heterogeneity and plasticity of cortical and spinal microglia, and suggest functional differentiation of microglia in different CNS compartments.

## Introduction

The CNS-resident macrophages, microglia, play critical roles in CNS development, homeostasis and inflammation. During development, microglia participate in neural circuit formation by regulating specific processes such as axon outgrowth and synapse maturation and pruning^1,2^. In adults, microglia actively survey the microenvironment to maintain homeostasis^3^. In response to infection or injury, microglia are activated to remove pathogens, damaged cells and dysfunctional synapse, and facilitate tissue repair^4^.

The diverse functionality likely arises from different microglial subtypes with distinct morphologies and biological activities. For instance, non-phagocytic activated microglia appear bushy with short thick branches, undergo rapid proliferation, and secrete pro-inflammatory signalling molecules^5^, whereas phagocytic activated microglia show an ameboid shape, and can travel to injury sites to engulf cell debris^3,5^. From the perspective of inflammation, activated microglia are classically categorized into pro-inflammatory M1 and anti-inflammatory M2 subtypes. M1 microglia produce pro-inflammatory mediators such as TNF-a, IL-1β, and NO^6,7^, while M2 microglia express anti-inflammatory mediators^8^. The biological basis of microglial heterogeneity is still poorly understood.

Single cell RNA-seq (scRNA-seq) provides powerful means to characterize microglial heterogeneity. Transcriptomic data generated from this approach revealed novel microglial subtypes^9^,^10^. Microglia appear more heterogeneous during early development, compared with adult stages^10,11^. Yet, microglia in adult brains are still plastic and can adapt to different phenotypes in response to pathogenic insults, including subtypes beyond the M1 and M2 categories. At least four activated microglial subtypes with hallmarks of inflammation, proliferation, interferon response, and demyelination respectively have been identified in mouse models of aging and neurodegenerative diseases^9–14^.

Emerging scRNA-seq evidence suggests spatial and temporal heterogeneity of microglia in the brain^3^. These findings indicate regional- and age-dependent plasticity of microglia. However, the mechanism and biological significance of the spatial- and temporal-regulated microglial heterogeneity are unclear.

In this study, we performed scRNA-seq analysis on unsorted cells dissociated from brain cortices and spinal cords to compare microglial heterogeneity in the cortex and the spinal cord of adult mice. We found that in Wt mice both cortical and spinal microglia consisted of homeostatic and inflammatory subtypes although with differential proportions, and that cortical and spinal microglia responded differently to age progression. We further demonstrated differential plasticity of microglia in cortices and spinal cords in a mouse model of HIV-associated neurological complications. The findings suggest a region-specific mechanism in controlling the expression of microglial phenotypes and the differentiation of microglial functions in different CNS compartments.

## Results

### Microglial heterogeneity in the brain cortex and the spinal cord from 2-month wild type adult mice

To understand the spatial heterogeneity of microglia in the CNS in an unbiased manner, we performed droplet-based scRNA-seq^15^ on cells dissociated from Wt adult mouse (2-month) brain cortices and spinal cords (Sup. Fig. 1a). We adapted and optimized a density gradient cell separation protocol^16^ to minimize cell death from dissociation processes and collected the fractions enriched with microglia for the downstream scRNA-seq processing (Sup. Fig. 1a). Cells dissociated from two cortices and two whole spinal cords were pooled respectively. Total of 19960 unsorted cells from cortices (10000) and spinal cords (9961) were sequenced to a depth of ~30,000 raw reads per cell. Cells with over representation of mitochondrial genes (>10%) or low number of detected gene (<300 genes/cell) were filtered out. 5211 cells from the cortex (52 % of cortical cells) and 3780 cells from the spinal cords (38% of spinal cells) passed quality control for downstream analysis. We performed t-distributed stochastic neighbor embedding (t-SNE) cluster analysis (Seurat v2.21) on cortical or spinal cells. The resolution 0.6 was used for all t-SNE analysis in this study. The t-SNE analysis identified 14 clusters (Fig. 1a; Sup. Fig. 1b; Sup. Fig. 1c), and individual clusters displayed characteristic expression of specific sets of genes (Sup. Fig. 1d) that defined the cell types of each cluster (Fig. 1a, Sup. Fig. 1c). We found all major types of cells in the clusters of cortical cells, including neurons, microglia, astrocytes and oligodendrocytes, with neurons markedly under represented (Fig. 1a). Similar cell types were detected in the spinal clusters, except that few neurons were present (Fig. 1b). These data indicate microglia were the predominant clusters on the t-SNE plots from both the cortical and spinal cells, with few neurons due to the fraction we collected (Sup. Fig. 1a).

**Figure 1.**
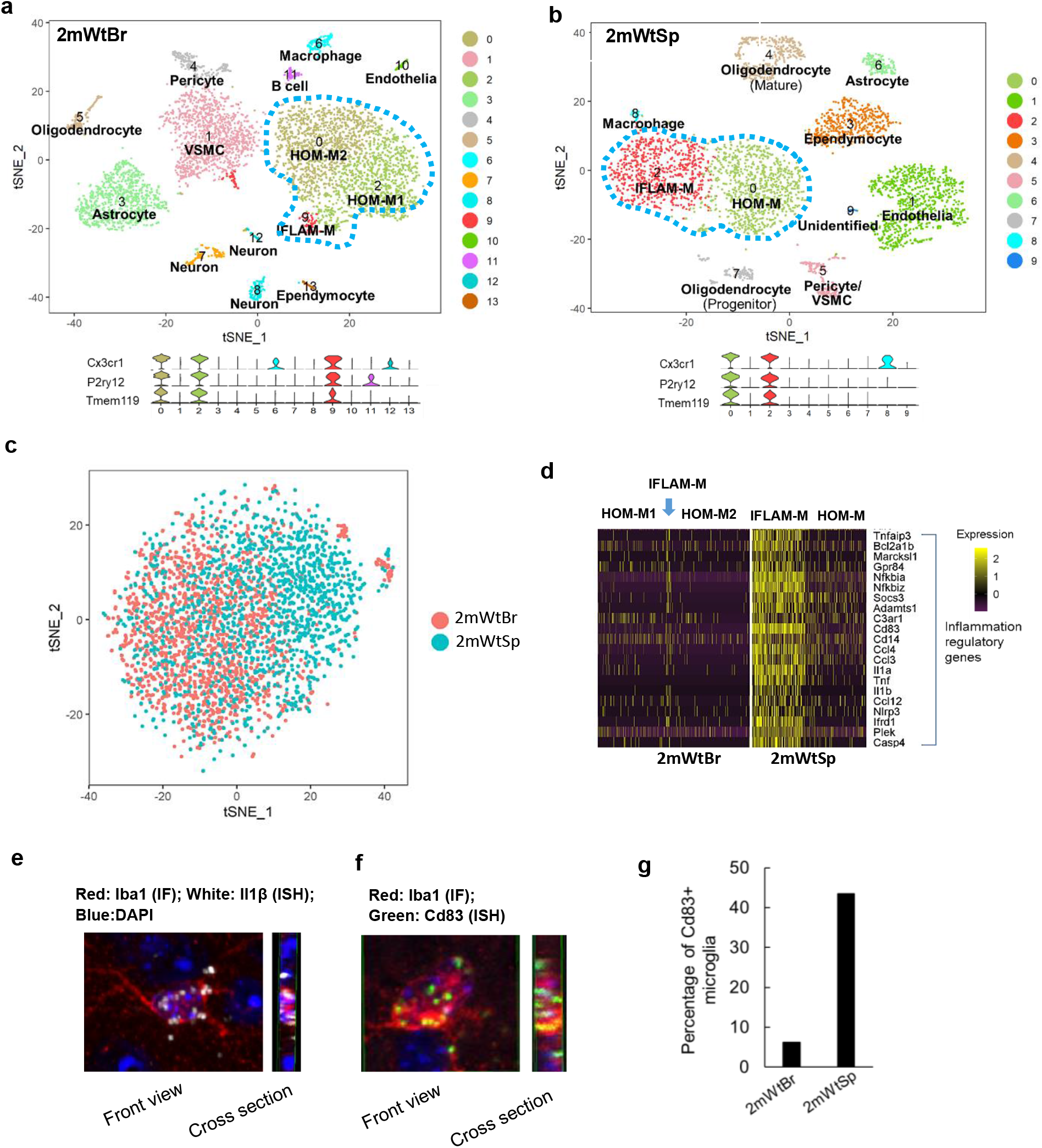
Single cell RNA-seq (scRNA-seq) analysis revealed region-specific microglial subtypes from the 2-month wild type (Wt) brain cortex (2mWtBr) and spinal cord (2mWtSp). **a** & **b.** t-SNE plots identified microglial clusters (blue dot line circled) in 2mWtBr (**a**) and 2mWtSp (**b**). Based on the expressed signature genes, the microglial clusters were defined as homeostatic microglia (HOM-M) and proinflammatory microglia (IFLAM-M). The cortical HOM-M was further segregated into HOM-M1 and HOM-M2 groups. Violin plots below t-SNE plots showed the expression of microglial markers Cx3cr1, P2ry12, and Tmem119 in t-SNE clusters. We defined the clusters with detectable expression of all three markers as microglia. VSMC: vascular smooth muscle cell. **c** Seurat Canonical Correlation Analysis (CCA) analysis showed that spinal microglia partially overlapped with cortical microglia, indicating cortical- and spinal-specific microglial heterogeneity. **d.** Gene expression heatmap showed the population of inflammatory microglia was big in spinal cord but small in the cortex. Each row: a gene; each column: a cell. **e**. A Z-stack confocal image of a cortical microglial cell (identified by Iba1 immunofluorescent staining (IF) with IL-1 β expression (revealed by RNAscope in situ hybridization (ISH)). **f.** A Z-stack image of a spinal microglial cell (labeled Iba1 immunostaining) with the expression of Cd83 (ISH). **g**. Population sizes of Cd83^+^Iba1^+^ microglia in the cortices and the spinal cords (n=2).

We used multiple gene markers, including Cx3cr1, P2ry12 and Tmem119, to identify microglia (Fig.1a; Fig. 1b). Three microglial clusters were identified from the cortex on the t-SNE plot (Fig. 1a), while two microglial clusters from the spinal cord (Fig. 1b). These observations indicated a differentiation between cortical and spinal microglia. To further visualize this difference, we performed Seurat Canonical Correlation Analysis (CCA) analysis on pooled cortical and spinal microglia. As shown in Fig. 1c, cortical and spinal microglia only partially overlapped, showing the differential transcriptomic expression of the two microglial populations.

To gain more insights into the biological basis of the differences between cortical and spinal microglia, we sought to determine the microglial subtypes of individual clusters revealed by t-SNE analysis, based on their uniquely expressed genes. We found that two major microglial clusters in the cortical t-SNE plot (clusters 0 and 2 in Fig. 1a) showed high expression of homeostatic genes (e.g. Cx3cr1, Tmem119, P2ry12, Csf1r)^11^, and thus named them as homeostatic microglia (HOM-M) (Fig. 1a; Fig. 1d). The HOM-M subtype constituted 94% of cortical microglia. As shown by the clusters on the t-SNE plot (Fig. 1a) and gene expression heatmap (Fig. 1d), the cortical HOM-M consisted of two computationally polarized groups, HOM-M1 and HOM-M2. HOM-M2 has slightly lower expression of genes coding ribosomal proteins (e.g. Rps11, Rps21, Rpl26) and Rgs10, the gene coding a member of regulator of G protein signaling family^17^ (Fig. 2a). HOM-M1 microglia were enriched with the expression of genes in various molecular pathways that are implicated in maintaining homeostatic functions such as P2ry12, consistent with a role of HOM-M microglia in damage-sensing^18^. We also found HOM-M in the spinal microglia, corresponding to cluster 0 on the spinal t-SNE plot (Fig. 1b). The spinal HOM-M constituted 55% of the microglial population. In contrast to the polarization of cortical HOM-M into two groups (HOM-M1 and HOM-M2), the spinal HOM-M microglia were not further polarized within the subtype into different clusters (Fig. 1b; Fig. 1d).

**Figure 2.**
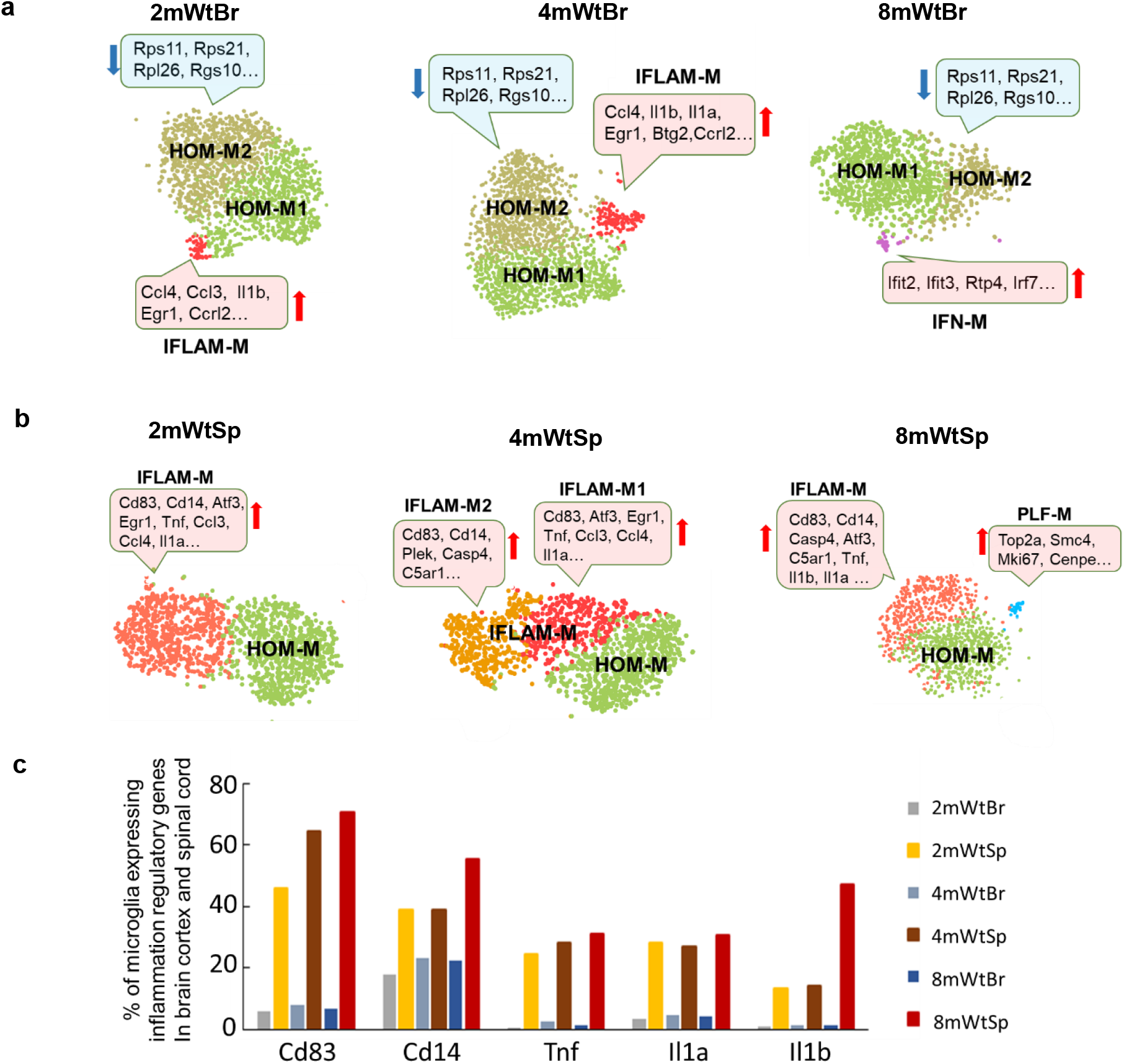
Different temporal plasticity of mouse cortical and spinal microglial heterogeneity. **a.** The temporal profiles of cortical microglial subtypes in 2-, 4-, and 8-month WT mice. HOM-M were polarized into HOM-M1 and HOM-M2 at all ages. HOM-M1 displayed with a characteristic down-regulation of genes including Rps11, Rps21, Rpl26, Rgs10 (blue boxes) genes coding for ribosomal protein and regulator of G protein signaling. IFLAM-M was a small subtype of cortical microglia identified at 2- and 4-month but not at 8-month. A subtype of interferon related microglia (IFN-M) appeared at 8-month. The signature high expressed genes in specific clusters were indicated in pink boxes. We only highlighted genes upregulated (red) or downregulated in the corresponding clusters (blue) compared to HOM-M1. **b**. The temporal profiles of spinal microglial subtypes in 2-, 4- and 8-month Wt mice. IFLAM-M were segregated into two sub-clusters (IFLAM-M1 and IFLAM-M2) at 4-month with their specific set of upregulated genes (pink boxes). At 8month, a small microglial cluster expressing genes regulating cell cycle and proliferation (PLF-M) appeared. **c.** Quantitative differences of cortical and spinal microglia expressing specific inflammation related genes, at different ages. Spinal microglia had a markedly higher portion to express a set of inflammatory regulatory genes than cortical microglia, at all ages checked.

Another microglial subtype was identified by their unique expression of genes regulating the immune response (e.g. Il1a, Il1b, Ccl3, Ccl4), and thus named inflammatory microglia (IFLAM-M). This subtype corresponded to cluster 9 in the cortical t-SNE plot (Fig. 1a). Microglia enriched with cytokine and chemokine genes (e.g. Il1b, Ccl2, Ccl4) were previously found from scRNA-seq analysis of human and mouse brains^11,14^. They were described as inflammatory microglia^14^ or preactivated microglia^11^. The IFLAM-M microglia were enriched with the expression of genes involved in cytokines such as Il1b (Fig. 1e), indicating the biological function of this subtype of microglia in mediating proinflammation. Cortical microglia from 2-month mice only had a small population (4%) of IFLAM-M (Fig. 1a; Fig. 1d). In contrast, IFLAM-M constituted 45% of spinal microglia (cluster 2 in Fig. 1b; Fig. 1d). The different sizes of cortical and spinal IFLAM-M populations were confirmed by the feature plots of microglial marker genes (Cx3cr1, P2ry12) and IFLAM-M enriched genes (Sup. Fig. 2), as well as by RNA in situ hybridization (RNAscope) of Cd83 (Fig. 1f; Fig.1g), one of the most consistent markers of IFLAM-M (Fig. 1d).

### The temporal differentiation of cortical and spinal microglial plasticity

The observations of differentiation of cortical but not spinal HOM-M and the drastic difference in population sizes of HOM-M and IFLAM-M in the cortex and the spinal cord suggest differential heterogeneity of cortical and spinal microglia in the 2-month mice. Next, we sought to determine if the cortical and spinal microglia showed different plasticity. To this end, we compared transcriptomes of cortical and spinal microglia at different animal ages (2-, 4- and 8-month). The remarkable difference in population sizes of HOM-M and IFLAM-M in the cortex and spinal cord revealed at two months of age were also seen at 4-, and 8-months (Fig.2a; Fig. 2b; Sup. Fig. 3). HOM-M were the majority cortical microglia, with 96%, 92% and 98% at 2-, 4 and 8-months respectively, whereas the cortical IFLAM-M only appeared at 2 and 4 months, with 4% and 8% respectively.

Same as the 2-month, cortical HOM-M were polarized into HOM-M1 and HOM-M2 at 4- and 8-months (Fig. 2a). Compared to HOM-M1, HOM-M2 expressed relatively lower levels of genes coding ribosomal protein and regulator of G protein signaling (e.g. Rps11, Rps21, Rpl26, Rgs10). The functional difference between these two clusters is unknown. However, the recent findings of microglial RGS10 as an anti-inflammatory and neuroprotective mediator^19^ suggest HOM-M1 and HOM-M2 may play different roles in CNS inflammation and homeostasis maintenance. In contrast to the cortical microglia, no polarization of spinal HOM-M was observed at all ages (Fig. 2b).

The cortical IFLAM-M comprised 4% and 8% of total cortical microglia at 2- and 4-months respectively (Fig.2a). These microglia expressed inflammatory cytokines (e.g. Il1, Ccl3, CCL4). At 8 months, the cortical IFLAM-M was not detected. Unlike the small cortical IFLAM-M clusters, the spinal IFLAM-M comprised a large population (41% ~ 54%) of spinal microglia. Spinal IFLAM-M displayed evident alteration during aging. At 2- and 8-month, they formed one cluster (Fig. 2b). While at 4-month, IFLAM-M were segregated into two clusters, IFLAM-M1 and IFLAM-M2 (Fig. 2b). IFLAM-M1 expressed genes (e.g. Atf3, Zfp36, Jun, Ccl4, Ccl3, Tnf, Il1α) that regulate acute response and inflammation^20^,^21^, while IFLAM-M2 expressed genes (e.g. Cd14, Gpr84, C5ar1, C3ar1) that module microglia activation^22,23^ and complement response^24^. In addition, we found that the percentage of microglia expressing Cd83, Cd14 and Il1b was increased in the spinal cord at 8-month (Fig. 2c) but not in the brain cortex, suggesting a polarization of spinal microglia toward a pro-inflammatory phenotype at this stage^22,25^.

At 8 months, a small cortical microglial cluster (2% of cortical microglia) with the signature of interferon-response genes (e.g. Ifit2, Ifit3, and Ifi204) emerged (Fig. 2a), and was named as interferon response microglia (INF-M). Interferon-related microglia were also found in an Alzheimer’s disease mouse model^13^ and aged mouse brains^14^. INF-M and the previously identified interferon-related microglia^26^ shared the expression of multiple interferon-stimulated genes, although fewer members were identified in INF-M. IFN-M was not detected in spinal microglia at the same stage (Fig. 2b).

A small (3%) spinal-specific microglial cluster appeared at 8 months, with a signature expression of a set of genes implicated in proliferation and cell cycle control (e.g. Mki67, Top2a) (Fig. 2b). We named this cluster as proliferation related microglia (PLF-M). Microglia expressing proliferative genes were identified in the developing white matter as proliferative-region-associated microglia (PAM)^10^. However, we did not detect in PLF-M the expression of phagocytosis-related genes, which was a signature of PAM^10^. PLF-M was not detected in the cortical microglia.

### Cortical and spinal microglial heterogeneity in HIV-1 gp120 Transgenic mice at two months of age

The results described above revealed microglial cortical- and spinal-specific heterogeneity, and the significance of this compartment-specific heterogeneity is suggested by its age-related differential plasticity. Because microglia are CNS resident immune cells, we sought to further confirm the biological relevance of the compartment-specific microglial heterogeneity by determining their response to pathogenic challenges. To this end, we used the HIV-1 gp120 transgenic (Tg) mouse, which expresses secreted gp120 in astrocytes and simulates HIV-associated neurological pathologies^27^. We found that microglial heterogeneity was increased both in the cortex and the spinal cord of the Tg mice at two months of age (Fig. 3a; Fig. 3b), compared with the age-matched Wt animals (Fig. 1a; Fig. 1b). Five microglial clusters were identified in the 2-month Tg cortex (Fig. 3a). Based on the transcriptomic signature, these clusters included the HOM-M (cluster 0 and 1), IFLAM-M (cluster 3), IFN-M (cluster 8) and PLF-M (cluster 7) (Fig. 3a). Compared with that in age-matched Wt mice, two new subtypes PLF-M and INF-M were induced in the transgenic mice (Fig. 3c; Fig. 3b; Fig.3d), with each constituting ~4% of total cortical microglia.

**Figure 3.**
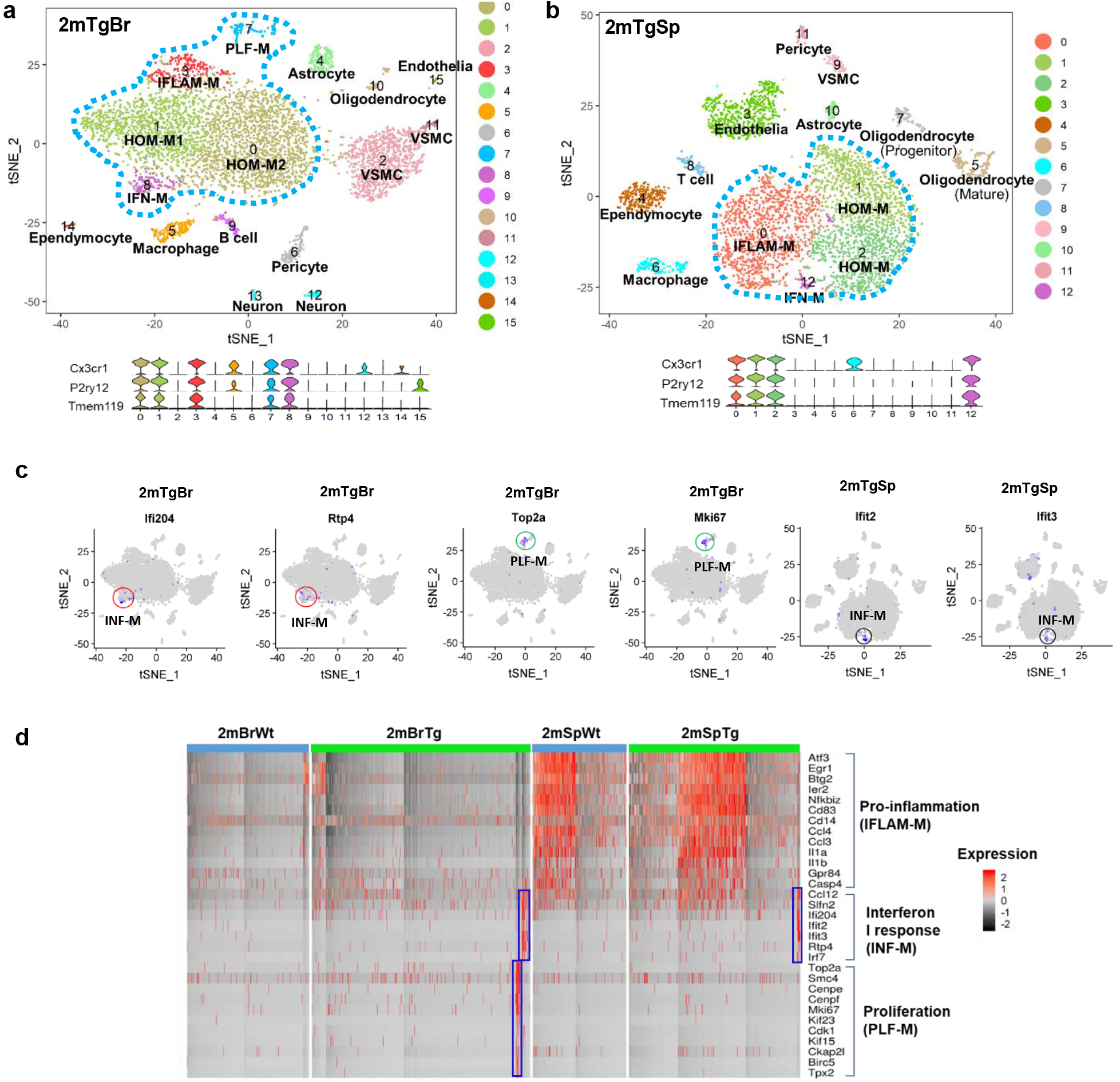
Region-specific microglial heterogeneity in 2-month HIV-1 gp120 transgenic (Tg) mice. **a** and **b**. t-SNE analysis identified five cortical (2mTgBr) and four spinal (2mTgSp) microglial clusters (circled by blue-dot lines) from the 2-month Tg mice. The microglial identity was defined by the co-expression of multiple microglial markers as shown in violin plots below t-SNE plots. **c.** Feature plots of signature genes of INF-M (red circles) and PLF-M (green circles) in 2mTgBr, and INF-M (black circles) in 2mTgSp demonstrate the portion of the corresponding subtypes in the cortical and spinal microglia. **d**. Heat map showed both INF-M and PLF-M microglia in the cortex but only INF-M in the spinal cord of the gp120 Tg mice. Gp120 did not significantly change the population of IFLAM-M in the cortex and spinal cord at 2-month. However, we can see the populations of cortical IFLAM-M are much larger in both Wt and Tg mice than in the Wt and Tg spinal cord, respectively.

Compared with cortical microglia, spinal microglia responded differently in the Tg mice at 2-months. Three spinal microglial subtypes were identified (Fig. 3b) including the HOM-M (cluster 1 and 2), IFLAM-M (cluster 0) and IFN-M (cluster 12) (Fig. 3b&3c). While the IFN-M subtype, which was not detected in WT cortical and spinal microglia at 2-months (Fig.1a; Fig.1b), was induced in both cortical and spinal microglial populations in the Tg mice at the same stage (Fig. 3d). The PLF-M subtype was only induced in the Tg cortical microglia but not in the spinal counterpart (Fig. 3d).

### Age-gp120 interaction differentially promoted cortical and spinal microglial heterogeneity

The data described above revealed differential plasticity of cortical and spinal microglial heterogeneity in response to age progression or gp120-induced pathogenesis. Next, we sought to test the interactive effect of age progression and gp120 on the expression of heterogeneity plasticity of cortical and spinal microglia. To this end, we determined the temporal profiles of cortical and spinal microglial subtypes at different ages (2-, 4-, and 8month) of the Tg mice. We observed that cortical microglia displayed distinct but overlapping subtypes as age progressed. As at 2 months, cortical microglial heterogeneity increased both at 4-month and 8-month in the Tg mice, compared with the cortical microglial subtypes from the age-matched WT mice (Fig.4a). Five cortical microglial subtypes were identified at 4-month Tg mice; they included HOM-M (including HOM-M1 and HOM-M2), IFLAM-M, INF-M and PLF-M, which were detected in the 2-month cortical microglia and a new subtype not detected in the 2-month cortical microglia (Fig. 4a; Fig. 4b). This new subtype comprised 11% of total cortical microglia, and was named as MYE-M because it had the signature expression of genes that promote demyelination and remyelination (e.g. Lpl, Cst7, Igf1, Spp1, Fabp5, Itgax)^28–30^. Compared to the age-matched Wt, the population of IFLAM-M increased in the 2 and 4-month Tg cortices (Fig. 4b; Fig. 8). The 8-month Tg cortical microglia contained overlapping clusters with 2- and 4-month Tg cortices except the MYE-M (Fig. 4a; Fig. 4b). MYE-M appeared in the 4-month cortex was not detected in the 8-month Tg cortex (Fig. 4a; Fig. 4b). In contrast, the INF-M and PLF-M remained (Fig. 4b). The population of INF-M gradually increased from 2- to 8-month in the Tg cortices and reached the highest at 8-month which covering 11% of the total cortical microglial population (Fig. 4b; Fig. 8). The population of PLF-M did not show significant changes in the Tg cortices along age progressing (Fig. 4b; Fig. 8).

**Figure 4.**
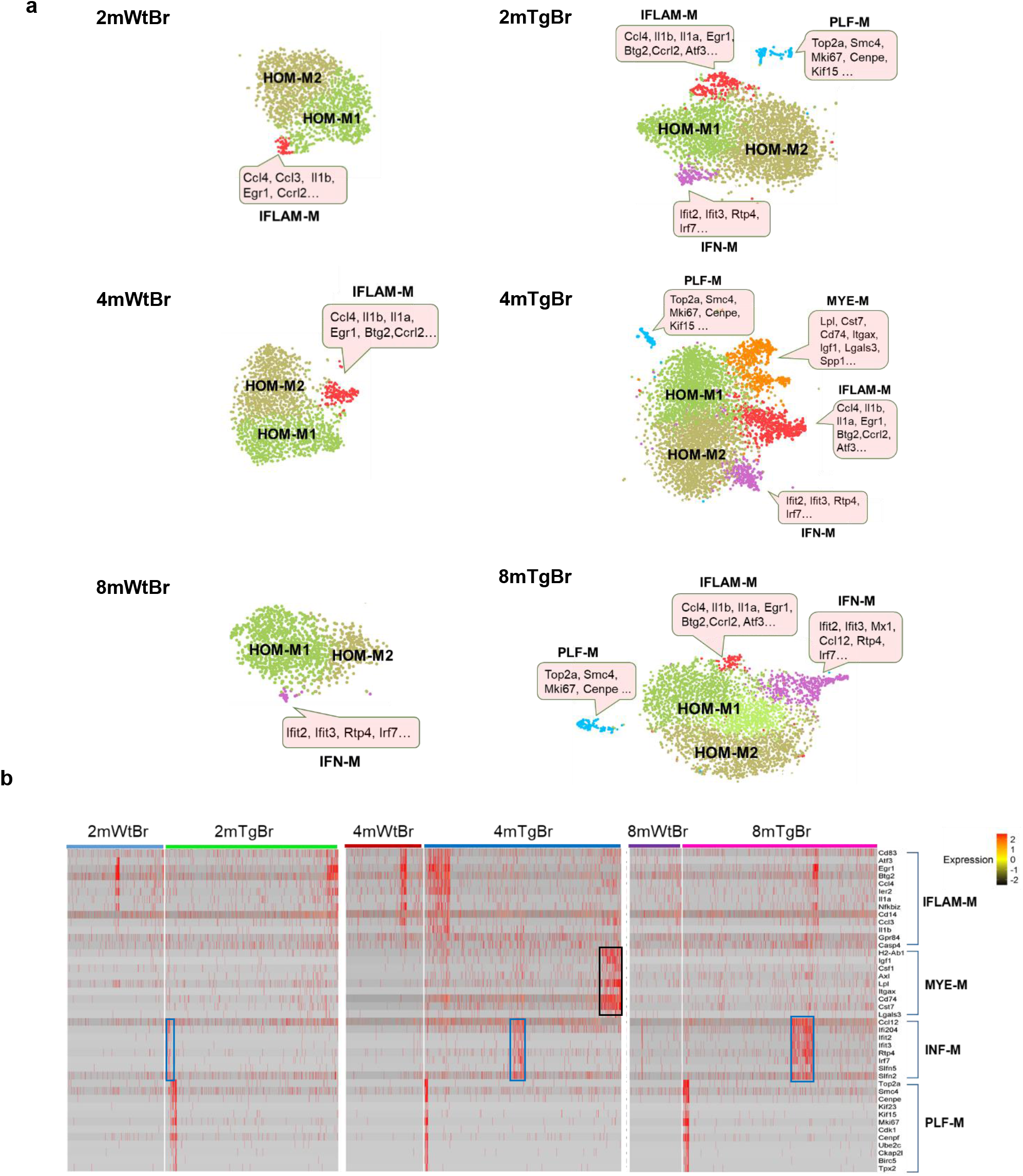
Comparison of the temporal profiles of cortical microglial heterogeneity of Wt and gp120 Tg mice. **a.** tSNE plots of Wt and Tg cortical microglia at different ages. The cortical microglial subtypes of the Tg mice at 2-, 4- and 8-month increased, compared to the Wt counterparts at the corresponding ages; this was particularly clear at 4-month. The signature genes expressed in the individual clusters were indicated in the pink boxes. **b**. Heatmap showed the expression of feature genes of IFLAM-M, MYE-M, INF-M and PLF-M subtypes. MYE-M (black rectangle) was only observed in 4-month Tg cortices, while INF-M (blue rectangle) gradually increased with age progression in Tg cortices.

We found MYE-M is particularly interesting as this type of microglia uniquely expressed a set of genes (e.g. Lpl, Cst7, Igf1, Itgax, Lgal3, Spp1) (Fig. 5a) which overlapped with the signature genes of the disease-associated microglia (DAM) identified from the brain of Alzheimer’s disease^9^. However, when comparing the upregulated genes in MYE-M (Supplementary Table 1 sheet 4TgBr_cluster_3_MYE-M) with that in DAM, we found that 15% (18/138) of the up-regulated genes in MYE-M overlapped with that from DAM (Fig. 5b). This observation suggests that MYE-M and DAM are related but distinct microglial subtypes induced in the HIV-1 gp120 transgenic and the Alzheimer’s disease models, respectively. To verify MYE-M *in vivo*, we performed *in situ* hybridization (RNAscope) of Cst7 mRNA and Lpl mRNA, in combination with Iba1 immunostaining. The results showed Cst7 mRNA and Lpl mRNA in clusters of microglia in the 4-month Tg cortex (Fig. 5c). We also performed immunostaining of Lgals3, another MYE-M marker. Lgals3^+^ Iba1^+^ microglia in the 4-month Wt cortex only consisted of 1~2% of cortical microglia. While Lgals3^+^ Iba1^+^ microglia comprised ~13% of the cortical microglia in the transgenic mice (Figure 5d, 5e), consistent with the finding of 11% MYE-M revealed by scRNA-seq analysis.

**Figure 5.**
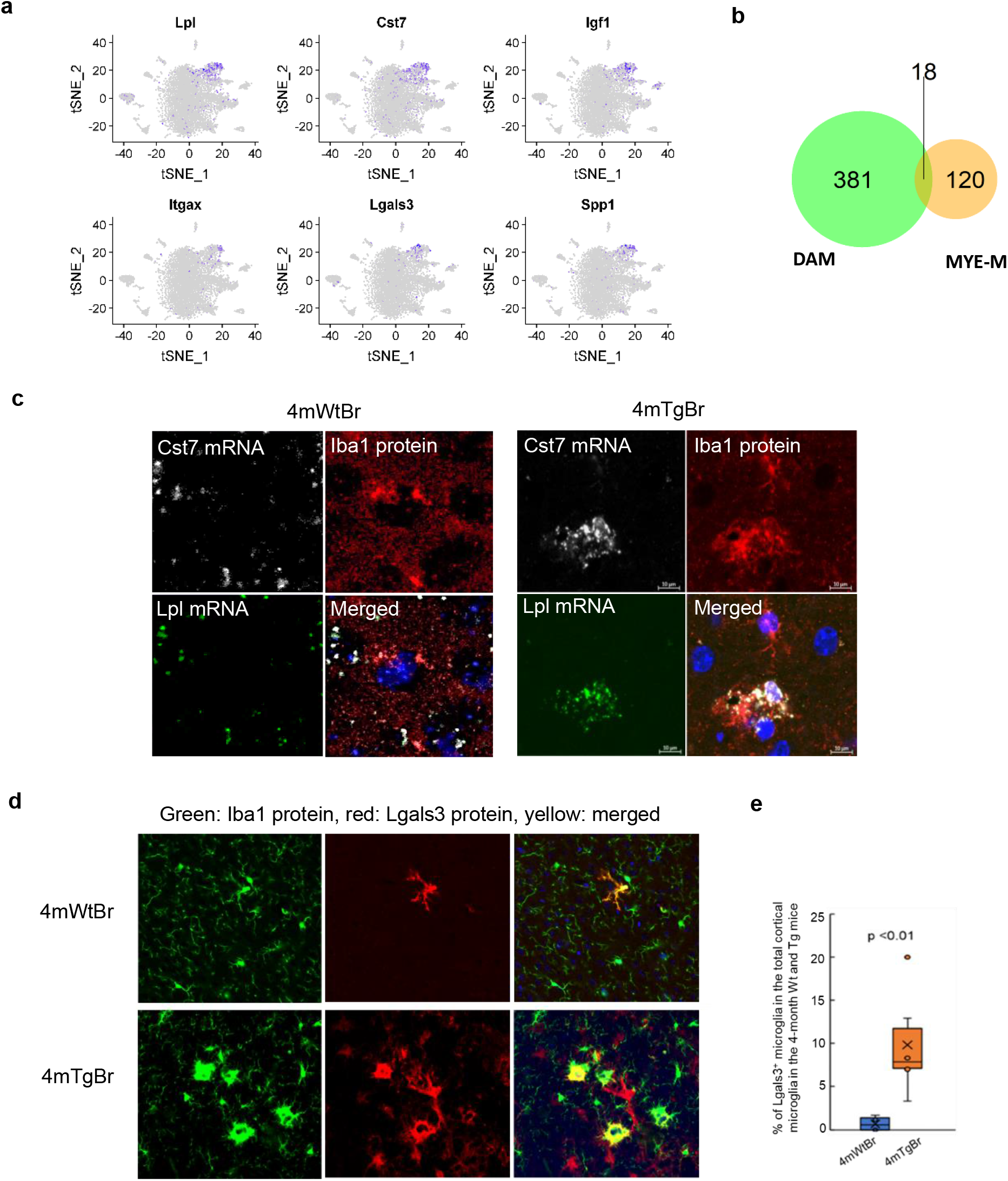
Verification of the MYE-M in cortical tissue at 4-month. **a.** Feature plots showing the signature genes of MYE-M in the 4-month Tg cortex (4mTgBr). **b.** Venn diagram of the genes upregulated in MYE-M and DAM (disease associated microglia) identified in Alzheimer’s disease. **c**. Cst7 and Lpl mRNAs (revealed by ISH) in cortical microglia (labeled by Iba1 immunostaining). Microglia with prominent Cst7 and Lpl expression were detected in Tg but not Wt cortices. **e.** Fluorescent immunostaining of the Lgals3^+^Iba1^+^ microglia in Wt and Tg cortices. **e**. Quantification of Lgals3^+^Iba1^+^ microglia in 4month Wt and Tg cortices.

In contrast to the differentiation of multiple new cortical microglial subtypes in the Tg mice at 2-, 4- and 8-month compared with the WT cortical microglia as described above, the spinal microglia in the Tg mice showed few new subtypes compared with their age-matched WT counterparts. We only observed the increase of microglial heterogeneity in the 2- and 4-month transgenic mice, with the appearance of INF-M (Fig.6a; Fig. 6b) that constituted ~2% of the total spinal microglia population. The INF-M was not detected in the 8-month Tg spinal cord of Wt or Tg mice (Fig. 6c). In addition, the PLF-M observed in the Wt spinal cord at 8-month was not detected in the 8-month Tg spinal cord (Fig. 6c). As in 4-month Wt spinal microglia, the IFLAM-M in Tg spinal microglia at the correspondent stage also differentiated into IFLAM-M1 and IFLAM-M2 groups (Fig. 6b).

**Figure 6.**
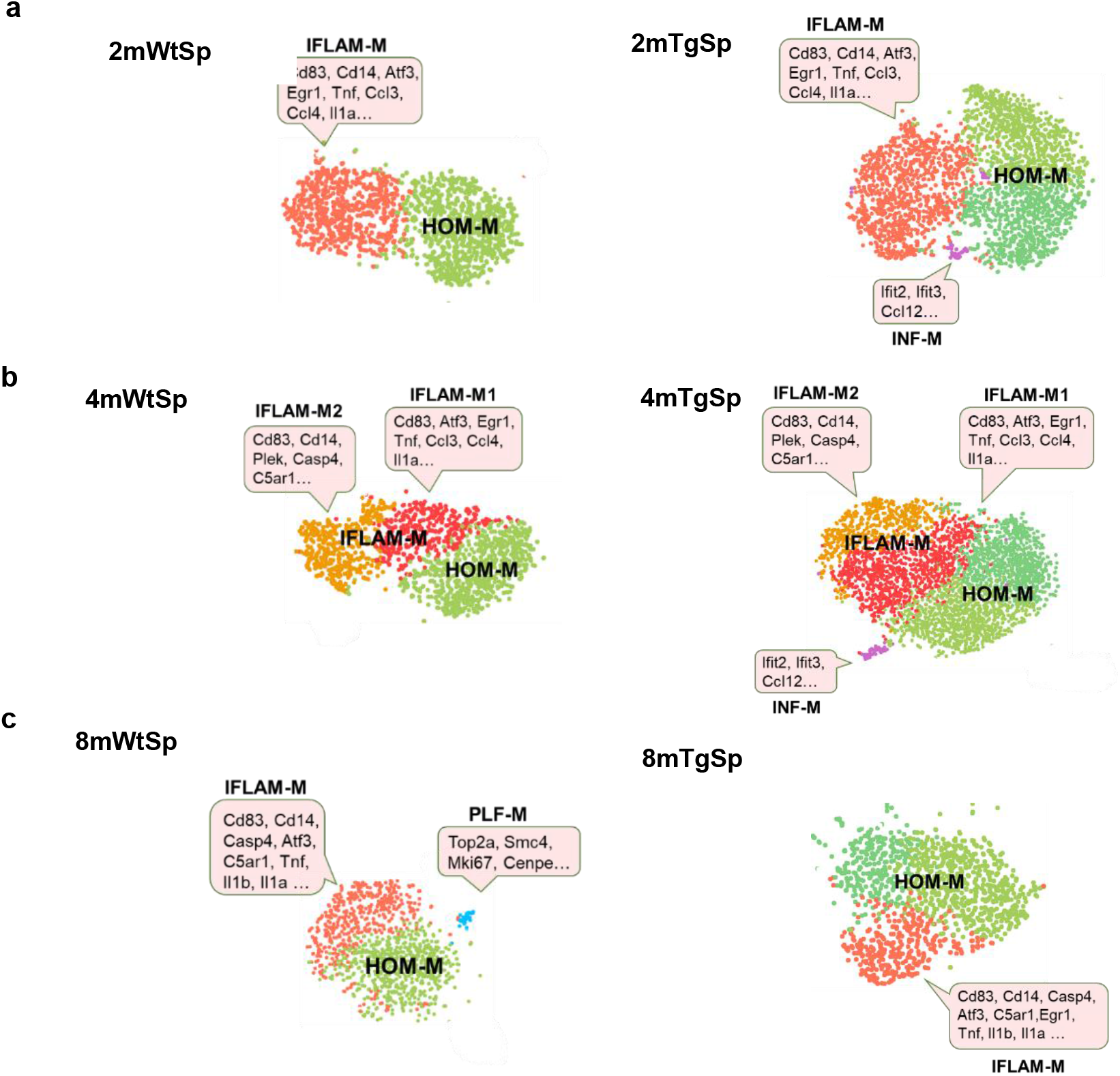
Comparison of the temporal profiles of spinal microglial heterogeneity of Wt and gp120 Tg mice. Compared to their Wt counterparts, INF-M was the only new subtype induced in 2- (**a**) and 4-month Tg mice (**b**). On the other hand, the PLF-M subtype detected in Wt mice at 8-month was not detected in the Tg mice at this age (**c**). The signature genes of specific clusters were indicated in pink boxes.

### Differential biological activity of cortical and spinal microglial subtypes in the gp120 transgenic mice

The above data revealed the differential plasticity of cortical and spinal microglial heterogeneity in the HIV-1 gp120 transgenic mouse. We next sought to investigate if the biological functions of the same microglial subtype in the cortex and the spinal cord were differentiated in the cortex and the spinal cord of the transgenic mice. To this end, we sought to compare the activity of the canonical pathways of cortical and spinal microglial subtypes in the WT and gp120 transgenic mice. Lists of up-regulated genes with the adjust p-value (<0.05) for individual microglial clusters (**Supplementary Table 1 and 2**) from 2-, 4-, 8-month Wt and Tg cortices and spinal cords were input into IPA to evaluate the change of individual biological pathways (e.g. immune response and nervous signalling pathways). We found many canonical pathways in the same subtype of the cortical and spinal microglia were differentially regulated in the Tg mice. For instance, some selected canonical pathways were evidently activated in both cortical and spinal HOM-M subtypes in WT mice, especially at 2- and 4-months (Fig. 7a, Fig7c). However, these pathways, except the neuroinflammation signalling, were severely attenuated in the cortical HOM-M of the transgenic mice across all ages (Fig. 7b). In the spinal cords, however, the attenuation of these pathways seemed to occur, but with to a lesser degree compared with their cortical counterparts (Fig. 7d).

**Figure 7.**
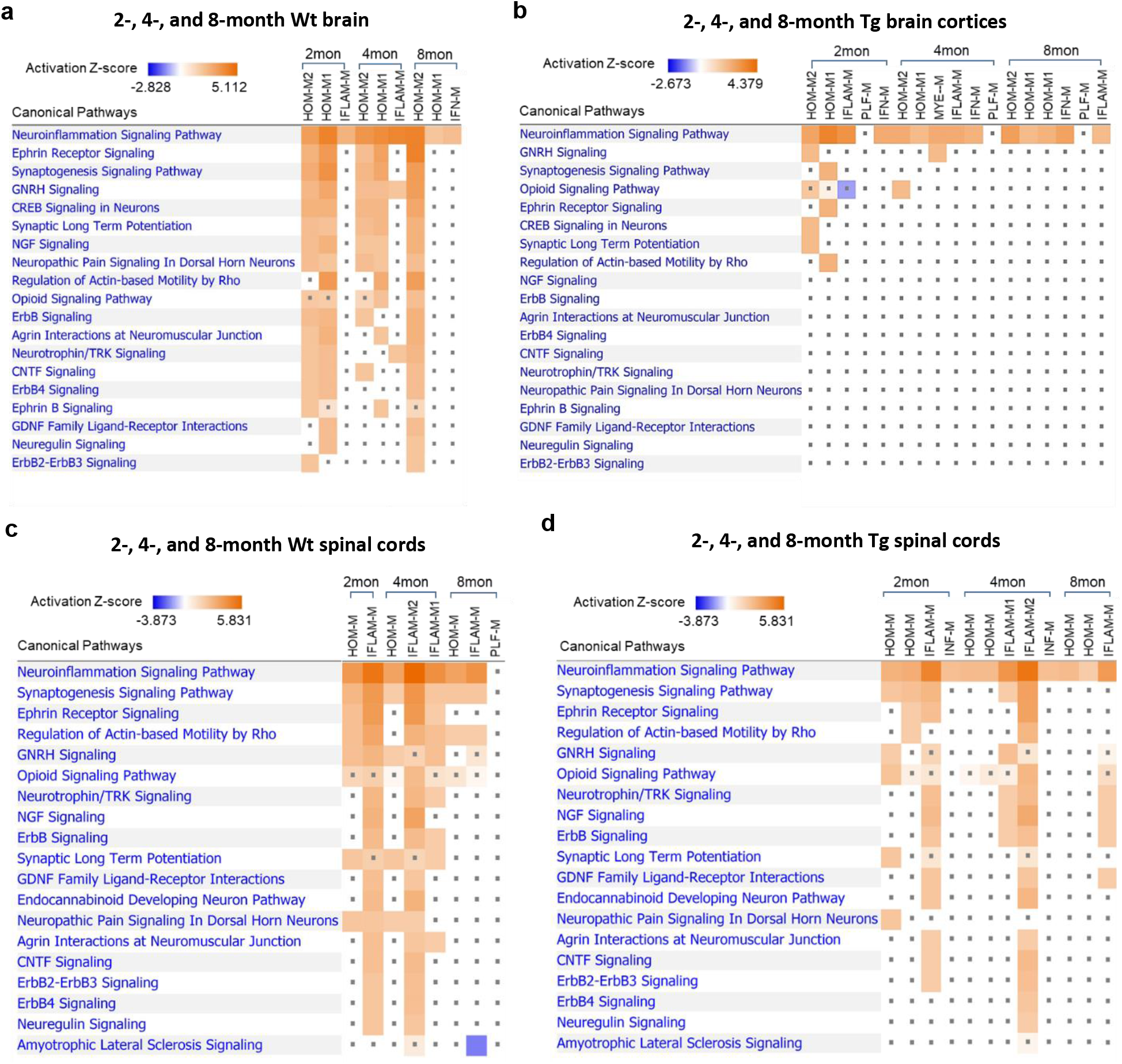
The effect of HIV-1 gp120 on cortical and spinal microglial nervous signaling pathways. **a**. Ingenuity Pathway Analysis (IPA) predicted the activated nervous signaling pathways of cortical microglia in the 2-, 4-, and 8-month Wt cortices. **b**. The activated cortical microglial signaling were severely suppressed by gp120 in the Tg brain cortices at all ages. **c**. IPA predicted the activated nervous signaling pathways of spinal microglia in the 2-, 4-, and 4-month Wt spinal cords. **d**. The activated nervous signaling of spinal microglia were differentially attenuated by gp120 in the Tg spinal cords. Note the signaling of spinal microglia were attenuated in a less degree compared to that of the cortical microglia. Orange: activated signaling. Blue: inhibited signaling. Dot: not significantly changed.

## Discussion

In this study, we compared microglia from the cortices and spinal cords using scRNA-seq approaches and characterized microglial subtypes in these CNS compartments based on clustering analysis. Notwithstanding the intrinsic technical caveats associated with scRNA-seq, the multi-way comparisons enabled by including samples from different tissues (cortices and spinal cords), different ages (2-, 4- and 8-month) and different mouse lines (Wt and gp120 Tg) allowed us to evaluate the consistency among the data sets. Our results reveal an overlapping but distinct microglial populations in the cortex and the spinal cord (Fig. 8). The differential heterogeneity is further shown by the differences in their plasticity under two neurophysiological conditions relevant to the biological function of microglia - life span progression and pathogen exposure. The findings indicate that microglia in the cortical and spinal compartments are adapted to the local environments to fulfil their unique biological functions, by differentiating distinct subtypes and/or expressing differential plasticity of the same subtypes.

**Figure 8.**
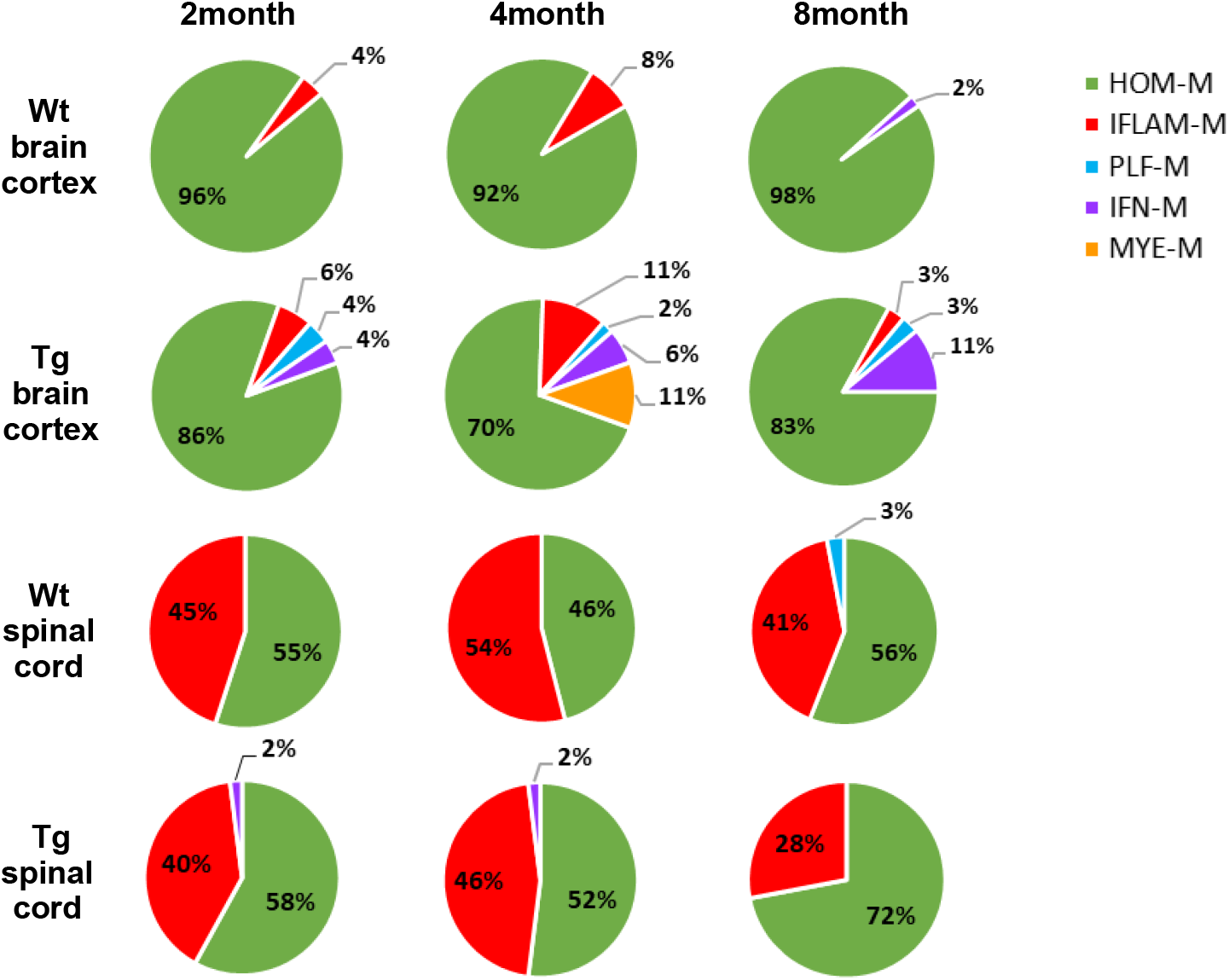
A summary pie charts show the population sizes of individual microglial subtypes in Wt and Tg cortices and spinal cords at 2-, 4-, 8-month.

### Cortical- and spinal-specific microglial heterogeneity

We found that microglia in the cortex and the spinal cord in 2- and 4-month WT mice consisted of two subtypes, HOM-M and IFLAM-M. However, the proportions between HOM-M and IFLAM-M were dramatically different in the cortex and the spinal cord. IFLAM-M comprised only a small portion (4-8%) of the cortical microglia, while in the spinal cord about half of microglia were IFLAM-M (Fig. 8). Because IFLAM-M express inflammatory genes and likely play a critical role in the inflammatory response, these observations indicate that the cortex may have a more limited capacity in microglia-mediated inflammatory response. In contrast, the large and more active population of the spinal IFLAM-M microglia suggest an increased capacity of microglia-mediated inflammatory response of the spinal cord. The plasticity of spinal microglia was also suggested by the observation the spinal-specific plastic changes of the IFLAM-M during age progression. While spinal IFLAM-M were detected as one cluster at 2-month, then differentiated into IFLAM-M1 and IFLAM-2 at 4-month, and returned to one cluster at 8-month (Fig. 2b). IFLAM-M1 expressed multiple signature genes of the classic M1 microglia (e.g. Tnf, Il1a, Il1b), and IFLAM-M2 expressed genes regulating cell activation and complementary response (e.g. Cd14, C5ar1, C3ar1).

In the cortex of WT mice, the IFLAM-M subset of microglia disappeared at 8-month (Fig. 8). Instead, a small cluster of IFN-M microglia was identified at this stage. INF-M microglia were of characterized with interferon-response genes, which are implicated in age-related modulation of cognitive function^31^. Given the reported aging-induced interferon-related microglia^13,14^, the observation of INF-M at 8-month may suggest interferon-related microglial activation during aging. It is unclear if there is a causal relationship between the disappearance of IFLAM-M and the appearance of INF-M in the 8-month cortex.

In the spinal cord at this stage, however, IFN-M was not observed (Fig. 8). Instead, a small cluster of PLF-M microglia appeared, constituting 3% of spinal microglia (Fig. 8). Because PLF-M microglia expressed genes regulating cell cycle and proliferation and were not observed in the cortex of WT mice at this stage, the appearance of spinal PLF-M indicated the occurrence of spinal-specific microglial proliferation during this age.

Together, the temporal comparative scRNA-seq analysis of cortical and spinal microglia of WT mice reveals that microglia in these two CNS compartments have distinct heterogeneity and age-related plasticity. These findings suggest compartment-specific differentiation of the biological function of microglia to maintain local homeostatic function of the CNS.

### Cortical- and spinal-specific plasticity of microglial heterogeneity in the HIV-1 gp120 transgenic mice

Temporal comparative scRNA-seq analysis revealed marked differences of microglial heterogeneity in the cortex and the spinal cord of the gp120 transgenic mice (Fig. 8). Compared to their Wt counterparts at different stages, in addition to the HOM-M and IFLAM-M subtypes, cortical microglia in the Tg mice formed three new subtypes, including INF-M, MYE-M, and PLF-M, while spinal microglia from transgenic mice only differentiated one new subtype, INF-M (Fig. 8). PLF-M and MYE-M were cortical-specific in the transgenic mice.

Although INF-M was shared by both the cortical and the spinal microglia, it showed different temporal profiles in these compartments. Cortical IFN-M were present at all three time points, and increasedin its proportions as age progressed (Fig. 8). On the other hand, spinal INF-M appeared at both 2- and 4-months with a small, steady percentage but disappeared at 8-months (Fig. 8). As type I interferon-induced reactive microglia were implicated in engulfing neuronal and synaptic elements^32^, the increase of the cortical INF-M population as age progresses may help clean damaged neural cells or debris induced by gp120. Type I interferon in the spinal cords is implicated in pain suppression^33^. It would be of interest for future studies to determine if INF-M contributes to the pathogenesis in the spinal pain neural circuits induced by HIV-1 neurotoxins.

PLF-M was a cortical-specific microglial subtype that presented at all stages in the transgenic mice but was absent in the spinal cord. The presence of PLF-M indicated that cortical microglia were re-entering cell cycles and proliferating in the transgenic mice. In contrast, the lack of PLF-M in the spinal cord suggested low proliferation of spinal microglia in the transgenic HIV model.

MYE-M was another cortical-specific subtype identified in the transgenic mice. However, unlike the PLF-M subtype, MYE-M was only detected in the 4-month Tg cortex (Fig. 8). Signature genes of MYE-M (e.g. Lpl, Cst7, Igf1, Itgax, Lgals3, Spp1) (Fig. 5a) were also found to be expressed in microglial subtypes (e.g. DAM) in neurodegeneration models^9,11,13,14^. We compared the upregulated genes from MYE-M and DAM and found only 18 out of 138 MYE-M enriched genes overlapped with that from DAM (Fig. 5b). These overlapping genes included Cst7, Lpl, and Itgax that are involved in myelinogenic processes^34^ and phagocytosis^35^. Combined RNA scope in situ hybridization with immunohistochemistry confirmed the gp120-induced MYE-M in the Tg brain cortices (Fig. 5c; Fig. 5d; Fig. 5e). DAM was implicated in clearance of damaged neurons associated with Alzheimer’s disease^9^, and it is tempting to speculate a similar role of MYE-M. However, the large portion of non-overlapping genes enriched in MYE-M and DAM suggests that these microglial subtypes are very different in their biological activities.

In summary, the findings from this study show the differential microglial heterogeneity in the mouse cortex and the spinal cord. They also demonstrate that microglia in the two CNS compartments have different plasticity in response to advancing age advancing and gp120-induced pathogenesis. It would be interesting for future studies to elucidate the potential intrinsic and/or extrinsic mechanisms by which the compartment-associated microglial heterogeneity and plasticity origin develop.

## Methods

### Animals

Gp120 Tg mice (from Dr. Marcus Kaul, Sanford-Burnham Medical Research Institute) express HIV-1 LAV gp120 under the control of the glial fibrillary acidic protein (GFAP) promoter^36^. All animal procedures were performed according to protocol 0904031B approved by the Institutional Animal Care and Use Committee at the University of Texas Medical Branch, and all methods were performed in accordance with the relevant guidelines and regulations.

### Cell dissociation from the brain cortices and spinal cords

To isolate cells for scRNA-seq, Wt and gp120 Tg mice were euthanized with 14% Urethane followed by direct decapitation directly. The brain cortices and whole spinal cords from the Wt and Tg mice of 2-, 4-, 8-month old (n=2, mixed gender) were rapidly dissected out respectively, and temporarily stored in 6ml cold Hibernate A/B27 (HABG) medium. HABG was prepared by adding 30 ml of Hibernate A (BrainBits, cat. no. HA), 600μl of B27 (ThermoFisher, cat. no.17504044), 88 μl of 0.5mM Glutamax (Invitrogen, cat. no. 35050-061), Penicillin-Streptomycin (ThermoFisher, cat. no.15070063) to final concentration of 2%, and DNase I (ThermoFisher, cat. no. AM2224) to final concentration of 80 U/ml. After completion of dissection, the cortical or spinal tissues were separately placed in 100 mm petri dishes on ice, cut into 0.5 mm pieces with a scalpel, and transferred to a 6 well culture plate containing 6ml papain digestive medium for each sample. Digestive medium was freshly prepared by dissolving 12 mg papain powder (Worthington, cat. no. LS003119) in 6 ml hibernate without calcium (BrainBits, cat. no. HACA) with 15 μl 0.5mM Glutamax, and DNase I at final concentration of 80 U/m. The digestive medium was activated at 37 °C for 20–30 min. Tissues were digested in the 37°C incubator for 1hr (agitating every 20 min), and then transferred with the digestive medium into a 15 ml corning tube on ice, followed by addition of 6ml cold HABG medium. The tissues were subsequently slowly triturated on ice with a 2ml Pasteur Pipet for 10–15 times. Then triturated cells 10–15 times by a fire-polished narrow-bored glass pipette into single cells. The dissociated cell suspension was passed through a 70μm strainer (MACS, cat. no. 130-098-462), and span at 400g for 5min at 4°C. After discarding the supernatant, the cell pellets were gently re-suspended in 6ml cold HABG.

### Separation of cells by density gradient centrifugation

Microglia in the cell suspension were enriched according to a published procedure^16^. Briefly, we diluted and prepared the OptiPrep (Sigma, cat. no. D1556) 4ml density gradient in a 15ml corning tube following the published protocol^16^. 6ml of the cell suspension was carefully pipetted onto the top of the density gradient and centrifuged at 800g (Beckman, Cs-16R Centrifuge) for 30min at 22°C (with acceleration and deceleration rates set at 0). The top layer containing cell debris was carefully aspirated. Fraction 1 enriched with oligodendrocytes, and fractions 2 and 3 enriched with neurons (Sup. Fig.1)^16^. The bottom fraction (~500μl) enriched with microglia^16^ was collected, mixed with 5ml cold HABG medium and centrifuged for 10min at 400g (Beckman, Cs-16R Centrifuge) at 4°C. The cell pellet was gently re-suspended in 3100 μl cold D-PBS and processed for additional removal of myelin and cell debris with Debris Removal Solution (Miltenyi Biotec, cat. no. 130-109-398), following manufacturer’s instructions. Cells were suspended in HABG medium. The viability was checked by trypan blue staining. Cell preps with viability >95% were shipped (overnight on ice) to Cincinnati Children’s Hospital Medical Center (to Dr. Potter) for Droplet-based scRNA-Seq.

### Droplet-based single-cell RNA-Seq

The droplet-based scRNA-Seq was based on a protocol from Macosko et. al (http://mccarrolllab.org/wp-content/uploads/2015/05/Online-Dropseq-Protocol-v.-3.1-Dec-2015.pdf). Hydrophobic-treated microfluidic devices were ordered from Nanoshift (Richmond, CA). 1.5 mL of cells (120k cells/ml) were co-flown through the device along with 1.5 mL barcoded beads (Chemgenes, 175k beads/ml) and droplet generation mineral oil (QX200, Bio-Rad Laboratories). Resulting emulsion was collected in a 50 ml tube at room temperature; subsequently, the emulsion was allowed to hybridize on ice for 45 min prior to droplet breakage. The excess oil was removed, and the emulsion was transferred to a 50 mL glass conical. 40 mL of cold 6X SSC was added to the emulsion, along with 1 ml perfluoro octanol with vigorous shaking to break droplets. Following droplet breakage, beads were collected from the supernatant into two 50 ml conical tubes. The tubes were spun at 1200 g for 5 min, pelleting the beads. The supernatant was subsequently removed, and the beads were transferred to a 1.5 ml low-adhesion tube. Following this the beads were rinsed once with 6x SSC, once with 5X RT buffer, and once with 1X RT buffer. Captured mRNAs hybridized to the beads were then reverse transcribed for 30 min at room temp. followed by 1.5 hr. at 42 °C with rotation in RT mix with 2000 U Maxima H Minus Reverse Transcriptase (ThermoFisher, MA) in 200 μl total volume. Reverse transcription was followed by an exonuclease treatment for 45 min to remove unextended bead primers. The nucleic acid on the beads were then PCR amplified (four cycles at 65 °C and 12 cycles at 67 °C annealing temperature with 3000 beads per 50 μl reaction). The cDNA from the PCR reaction was purified using 0.7x volume of SPRIselect beads (Beckman Coulter, cat. no. B23318). The quantity and quality of cDNA was measured using an Agilent Bioanalyzer hsDNA chip. To generate a library cDNA was fragmented and amplified (12 cycles) using the Nextera XT DNA Sample prep kit with three separate reactions of 600, 1200 and 1800 pg input cDNA. The libraries were pooled and purified twice using 0.7X volume of SPRIselect beads. The purified libraries were quantified using an hsDNA chip and were sequenced on the Illumina HiSeq 2500 using the sequencing parameters described in the DropSeq protocol. The libraries were run on a single lane targeting 150 million reads per library.

### Computational methods

Drop-seq FASTQ files were processed using the standard pipeline (Drop-seq tools version 1.12 from McCarroll lab, http://mccarrolllab.org/wp-content/uploads/2016/03/Drop-seqAlignmentCookbookv1.2Jan2016.pdf). Generated read counts were analysed in R version 3.4.4 (R Core Team 2018) via Seurat 2.2.1^37^. following its standard pre-processing and clustering workflow (https://satijalab.org/seurat/v3.1/pbmc3k_tutorial.html). Briefly, from the distribution of reads, the data was filtered to a minimum of 300 cells per gene and 30 genes per cell. This was followed by UMI and mitochondrial filtering and normalization prior to selecting all highly variable genes falling within a selected cut-off window for PCA clustering. Statistically significant PCs (p-value < 0.001) were used in cluster determination to produce heatmaps and t-SNE plots at a resolution of 0.6. Clusters were annotated based on the expression level of canonical marker genes and gene expression visualized using feature maps. Differential gene expression across these clusters and their cell types was carried out using a standard AUC classifier, returning only genes with adjusted p-values < 0.05. These genes were then used to search for enriched pathways via IPA (QIAGEN Inc.). Seurat violin plots that were consolidated using graphical packages in R^38^,^39^. A number of helper functions were also employed ranging from the data input, manipulation and display^40–42^ to color schemes^43^.

### RNAscope in situ hybridization (ISH) and immunofluorescent (IF) staining

Mice were anesthetized with euthanized with 14% Urethane and transcardially perfused with 30ml cold 1xPBS. The cortex and the spinal cord were collected, and fixed in freshly prepared 4% PFA for 24h at 4°C. The tissues were then immersed in 30% sucrose in 1xPBS at 4°C until they sank to the bottom, embedded in OCT mounting medium, and stored in an air-tight container at −80°C prior to sectioning. The mounted tissue blocks were equilibrated to −20°C in a cryostat for ~1 hour prior to sectioning. 10−20 μm cryo-sections were cut, mounted onto SuperFrost® Plus slides, and stored in air-tighten slide boxes in zip-lock bags at −80°C until use.

Probes for genes Cd83 (cat. no. 416761), Il1b (cat. no. 316891), Cst7 (498711-C2), Lpl (cat. no. 402791) were purchased from Advanced Cell Diagnostics (Newark, CA). ISH procedures, including tissue pre-treatment and probes hybridization, were performed according to the protocol of RNAScope® Multiplex Fluorescent V2 Assay provided by the manufacturer (Advanced Cell Diagnostics). Immediately after the last wash of ISH, the sections were rinsed with 1xPBS (3×5min), followed by immunostaining of Iba-1 and nuclear staining by DAPI as described (Dual ISH-IHC, Advanced Cell Diagnostics). The stained sections on glass slides were mounted with ProLong Gold Antifade Mountant (Invitrogen, cat. no. P36930) and stored overnight at 4°C before imaging (Zeiss LSM 880 confocal microscope system). Anti-Iba1 antibody (abcam, cat. ab178847) was used at1:200 dilution.

### Canonical pathway analysis

To evaluate the effects of HIV-1 gp120 on microglial function, upregulated genes with adjusted p-value of < 0.05 from each cluster were imported into Ingenuity Pathway Analysis (IPA) (QIAGEN Inc.). IPA Canonical Pathways Analysis was performed to visualize the activity of individual pathways. The significance (p value) of the association between the data set and the specific canonical pathway was computed by Fisher’s exact test (https://chhe.research.ncsu.edu/wordpress/wp-content/uploads/2015/10/IPA-Data-Analysis-training-slides-2016_04.pdf) and Benjamini-Hochberg multiple testing correction, with p < 0.05 as the threshold of significance. The activation Z-score was used to identify the biological pathway that were activated or inactivated. An absolute z-score of≥2 was considered significant, with z-score ≥2 for activated pathways and a z-score < −2 for inactivated pathways.

