## Supplemental Figures for "Single cell RNA-seq analysis reveals compartment-specific heterogeneity and plasticity of microglia"

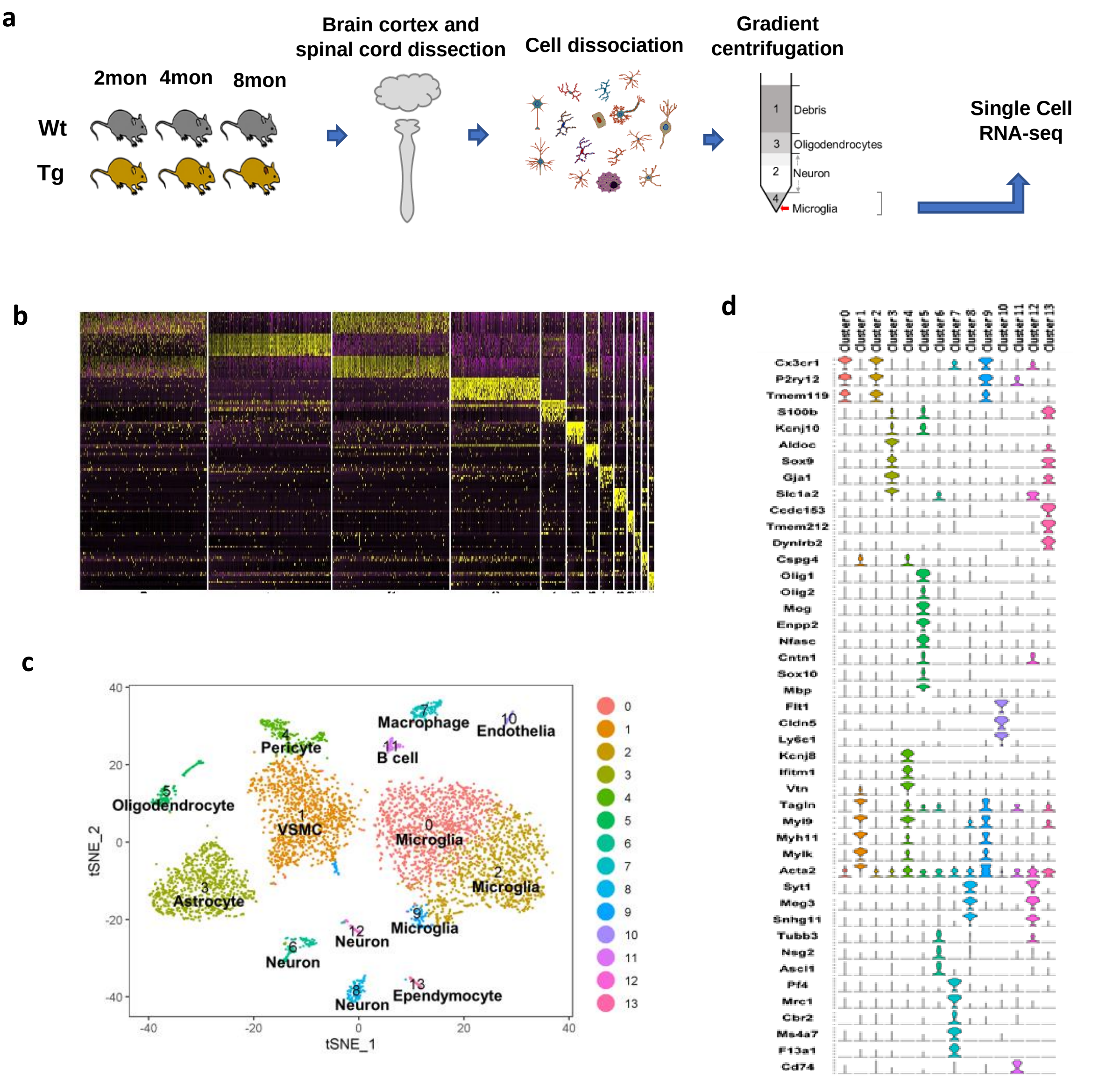


**Supplementary Figure 1. Workflow of the single cell RNA-seq (scRNA-seq) experiment and data analysis of cortical and spinal microglia of Wt and gp120 transgenic mice. a**: A diagram showing the workflow of cell dissociation and single cell RNA-seq (scRNA-seq)**.** Fraction 4 enriched with microglia was processed for the scRNA-seq. **b**. Expression heatmaps of the top 10 variable genes in each cluster demonstrated a well separated cluster. **c**. t-SNE plots displaying different cell types identified. VSMC: vascular smooth muscle cell. **d**: Combined Violent Plots of cell type-specific gene markers for individual clusters.

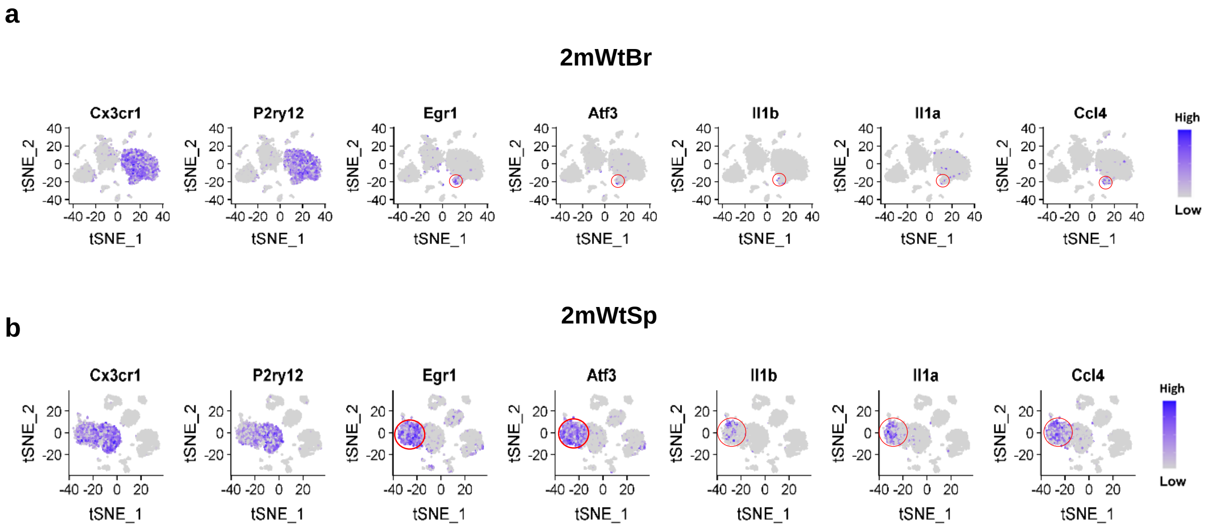


**Supplementary Figure 2. Feature plots showing the marked differences of IFLAM-M population sizes in the cortex (2mWtBr) and the spinal cord (2mWtSp) of Wt mice at 2-month.** Feature plots of pan-microglial marker genes Cx3cr1 and P2ry12 identifying the microglial populations in the t-SNE plots of cortical (a) and spinal (b) cells. Feature plots of IFLAM-M signature genes (Egr1, Atf3, Il1a, Il1b, Ccl4) showing the population of IFLAM-M (red circled) in the cortex (a) and spinal cord (b). Blue: high expression. Gray: low expression.

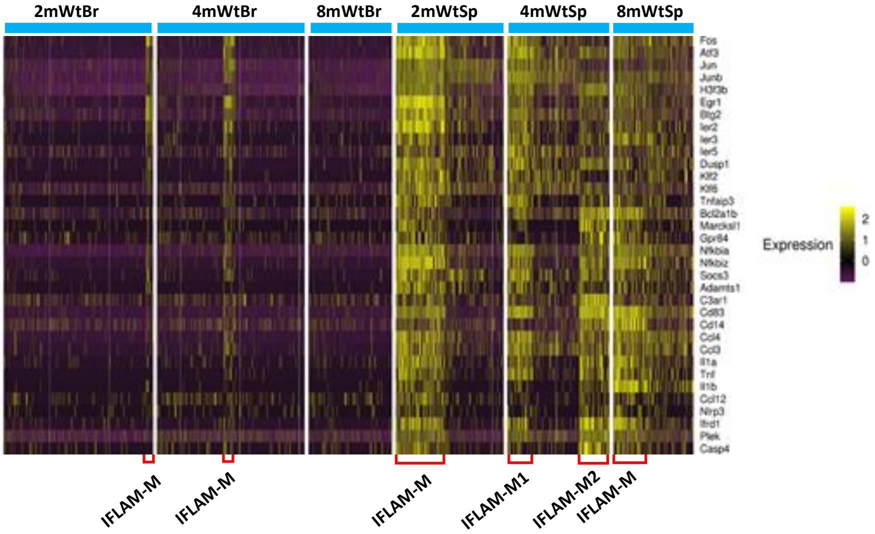


**Supplementary Figure 3. Gene expression heatmaps showing the IFLAM-M populations (red brackets) in the cortices and spinal cords of WT mice at 2, 4 and 8-month.**
